# Protocerebral bridge neurons that regulate sleep in *Drosophila melanogaster*

**DOI:** 10.1101/2020.10.30.361899

**Authors:** Jun Tomita, Gosuke Ban, Yoshiaki S. Kato, Kazuhiko Kume

**Affiliations:** Department of Neuropharmacology, Graduate School of Pharmaceutical Sciences, Nagoya City University, Nagoya 467-8603, Japan

## Abstract

The central complex is one of the major brain regions that control sleep in *Drosophila*, but the circuitry details of sleep regulation have yet to be elucidated. Here, we show a novel sleep-regulating neuronal circuit in the protocerebral bridge (PB) of the central complex. Activation of the PB interneurons labeled by the *R59E08-Gal4* and the PB columnar neurons in the *R52B10-Gal4* promoted sleep and wakefulness, respectively. A targeted GFP reconstitution across synaptic partners (t-GRASP) analysis demonstrated synaptic contacts between these two groups of sleep-regulating PB neurons. Furthermore, we found that activation of a pair of dopaminergic (DA) neurons projecting to the PB (T1 DA neurons) decreased sleep. The wake-promoting T1 DA neurons and the sleep-promoting PB interneurons formed close associations. *Dopamine 2-like receptor* (*Dop2R*) knockdown in the sleep-promoting PB interneurons increased sleep. These results indicated that the neuronal circuit in the PB regulated by dopamine signaling mediates sleep-wakefulness.

## Introduction

Sleep is a basic physiological state evolutionarily conserved among animal species (Allada and Siegel, 2008; Cirelli and Tononi, 2008). The fruit fly, *Drosophila melanogaster*, has emerged as a powerful model system for uncovering the molecular and cellular basis of sleep-wakefulness. Sleep in *Drosophila* is defined by behavioral features including circadian control of the timing of sleep, homeostasis of the amount of sleep and increased arousal threshold as in other animal species that do not exhibit the characteristic electroencephalogram patterns of sleep (Campbell and Tobler, 1984).

The central complex is a unique midline neuropil structures in the adult insect brain, composed of four substructures, the protocerebral bridge (PB), the fan-shaped body (FB), the ellipsoid body (EB) and the noduli (NO), and serves as a higher-order integrator for sensory information and various motor control (Homberg, 2008). Accumulating evidence from a number of neurogenetic studies has established a crucial role for the central complex in *Drosophila* sleep regulation. The sleep-promoting neurons having extensive presynaptic arborizations in the dorsal layer of the FB (dFB) has been identified (Donlea et al., 2011; Ueno et al., 2012). Dopamine is a neurotransmitter that is essential for maintaining wakefulness in *Drosophila*, similar to mammals (Andretic et al., 2005; Kume et al., 2005). The protocerebral posterior medial 3 (PPM3) cluster of dopaminergic (DA) neurons projecting to the dFB neurons controls sleep through Dopamine 1-like receptor 1 (Dop1R1) signaling (Liu et al., 2012; Ueno et al., 2012). Neurons that project to the ventral layer of the FB promote sleep and mediate consolidation of long-term memory (Dag et al., 2019). Homeostatic mechanism of sleep would be associated with changes in neuronal activity of the sleep-promoting dFB neurons and their functional upstream neurons, the R5 neurons in the EB (Donlea et al., 2014; Liu et al., 2016). In addition, the dFB neurons form a recurrent circuit with the EB R5 neurons via helicon cells to regulate sleep homeostasis (Donlea et al., 2018). Regarding circadian regulation of sleep-wakefulness, recent studies have shown that sleep-modulating dorsal neurons 1 (DN1) of circadian clock neurons and EB ring neuron subtypes are functionally connected via sleep-promoting tubercular-bulbar (TuBu) neurons and that this circuit is likely to regulate sleep-wakefulness (Guo et al., 2018; Lamaze et al., 2018). Besides, sleep-promoting lateral posterior neurons (LPNs), another group of clock neurons, appear to form close associations to the sleep-promoting dFB neurons and activate these neurons (Ni et al., 2019). Detailed mechanisms for regulating sleep-wakefulness by the central complex are being clarified. However, because the four substructures of the central complex are interconnected, a population of central complex neurons other than the FB neurons and the EB R5 neurons should also be implicated in the control of sleep-wakefulness.

Here, we begin to explore a novel sleep-regulating neuron in the central complex and discover that the sleep-promoting PB interneurons have direct neuronal connections to the wake-promoting PB columnar neurons projecting from the PB to the FB and the NO. Moreover, we reveal that T1 cluster of DA neurons regulates sleep by acting on the sleep-promoting PB interneurons through D2 dopamine receptor signaling.

## Results

### The PB defective mutant flies, *nob^KS49^* decreased sleep

In order to examine the role of the central complex in sleep regulation, we tested eleven mutant strains with morphological defects in the central complex. These mutant flies were isolated histologically following ethyl methanesulfonate mutagenesis, except for *agir^X1^*, which was isolated from a wild population (Strauss and Heisenberg, 1993). Flies were entrained to 12 h light : 12 h dark (LD) cycles for at least 3 days, then maintained in constant dark (DD) conditions. Locomotor activity was recorded in both LD and DD conditions. Sleep was defined as periods of immobile state lasting 5 min or longer, as previously described (Hendricks et al., 2000; Shaw et al., 2000; Huber et al., 2004; Kume et al., 2005). Because most of these structural mutants are still uncharacterized, their total daily activity and sleep were compared to those of three control strains (Canton-S (CS), *y w* and *w^1118^*) under DD conditions. There were no significant differences in total daily activity and sleep between the three control strains (*Figure 1A,B*). Among the central complex mutants tested, *nob^KS49^* flies showed significant hyperactivity compared with two control lines (CS and *y w*). *nob^KS49^* mutants have a defect in the central part of PB (Strauss et al., 1992). No difference was seen in waking activity index, defined as total daily activity divided by the active period length, between *nob^KS49^* and controls (*Figure 1C*), suggesting hyperactivity in *nob^KS49^* flies is not attributed to aberrant locomotion. On average, total daily activity significantly increased by about 2-fold in *nob^KS49^* relative to controls (*Figure 1A*). In contrast, total daily sleep of *nob^KS49^* flies decreased to approximately half of that of the CS and *y w* controls (*Figure 1B*). These results suggest that PB is involved in sleep regulation.

**Figure 1.**
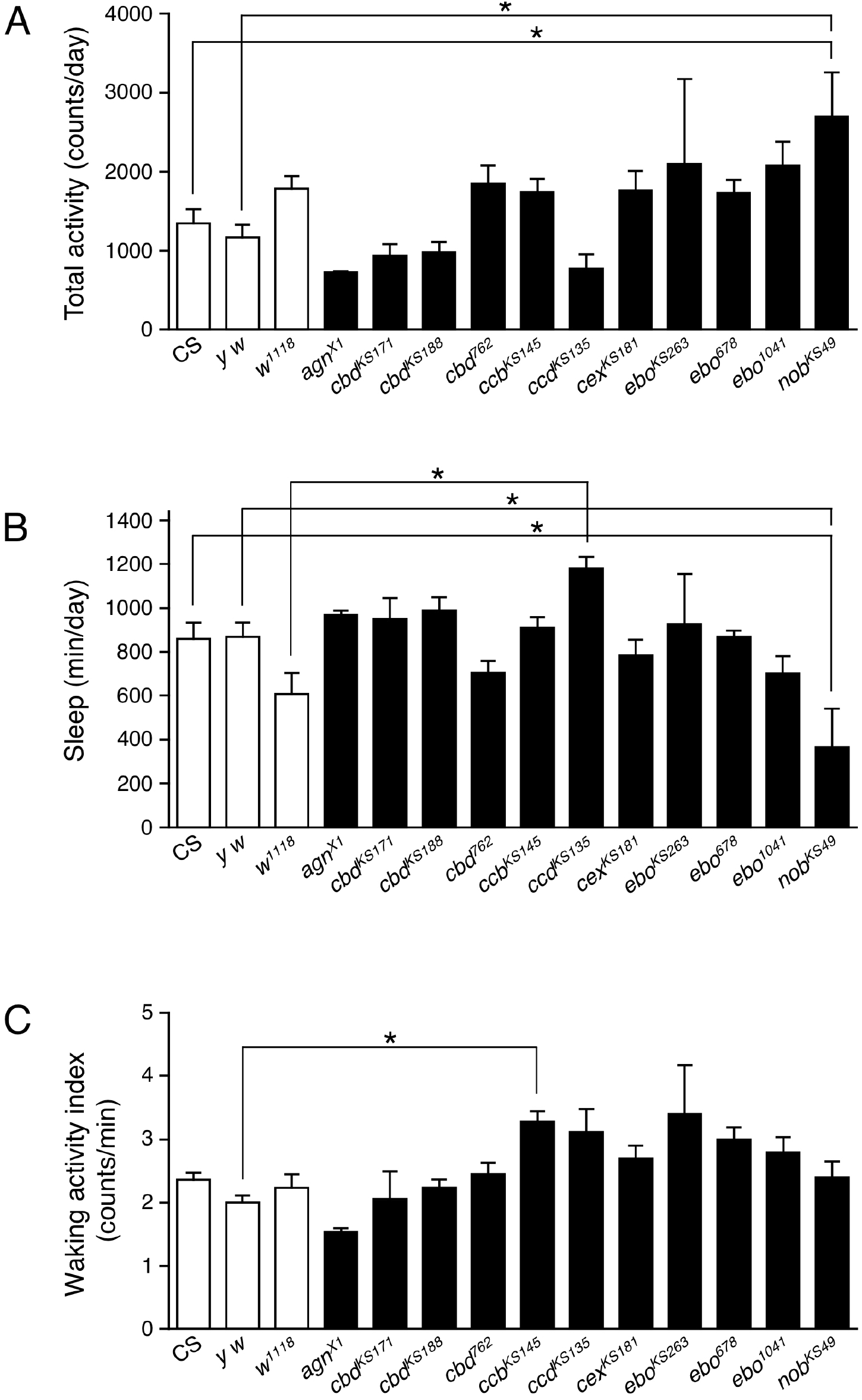
The PB defective *nob^KS49^* decreased sleep. Total daily activity **(A)**, total sleep **(B)** and waking activity index **(C)** for control flies (Canton-S, *y w* and *w^1118^*, white bars) and mutant strains with structural defects in the central complex (black bars) in constant dark (DD) conditions. Data are averaged for 3 days and are presented as mean ± standard error of the mean (SEM) (*n* = 3-11 for each group). Groups with asterisks indicate statistically significant differences (Tukey-Kramer HSD test for normally distributed data, *p* < 0.05).

### Activation of *Gal4*-expressing neurons in *Gal4* drivers that express in the PB affected sleep

To further explore the involvement of PB neurons in sleep regulation, we examined the effects of transient thermogenetic activation of a subset of PB neurons using the temperature-sensitive cation channel dTrpA1 on the amount of sleep. For this, fourteen *Gal4* driver lines that express strongly in PB neurons (*PB-Gal4*) were selected from the Gal4 image data of the FlyLight Image Database (https://flweb.janelia.org/cgi-bin/flew.cgi, Jenett et al., 2012). Progenies from the control (*PB-Gal4 × w^1118^*) and the experimental (*PB-Gal4 × UAS-dTrpA1*) crosses were grown at 22°C to adulthood. Sleep was measured in adult progeny flies in DD following LD cycles. These flies were transferred from 22°C to 29°C for 24 h to conditionally activate dTrpA1-expressing neurons and then returned to 22°C. Of the 14 *PB-Gal4* drivers tested, the acute activation of neurons with the 9 *Gal4* drivers significantly decreased sleep. In particular, the decrease in the sleep amount induced by activation of neurons using the *R52B10-Gal4* was most remarkable and flies hardly slept during neuronal activation as previously reported (*Figure 2A-C*) (Liu et al., 2016). Although clear homeostatic sleep recovery (sleep rebound) after sleep loss by the thermogenetic activation using the *R52B10-Gal4* was reported (Liu et al., 2016), such a clear sleep rebound was not observed in our experimental conditions (*Figure 2B,C*). On the other hand, the activation of Gal4 expressing neurons in the 3 of 14 *PB-Gal4* drivers significantly increased sleep (*Figure 2A*). The largest effect on sleep induction was caused by the neuronal activation using the *R59E08-Gal4*. In this *Gal4* driver, the amount of sleep in the experimental flies peaked in the middle of the subjective day at 29°C and was maintained at the peak level during neuronal activation (*Figure 2D*). Sleep amount in the experimental flies was significantly reduced compared with that in the control flies after stopping the activation.

**Figure 2.**
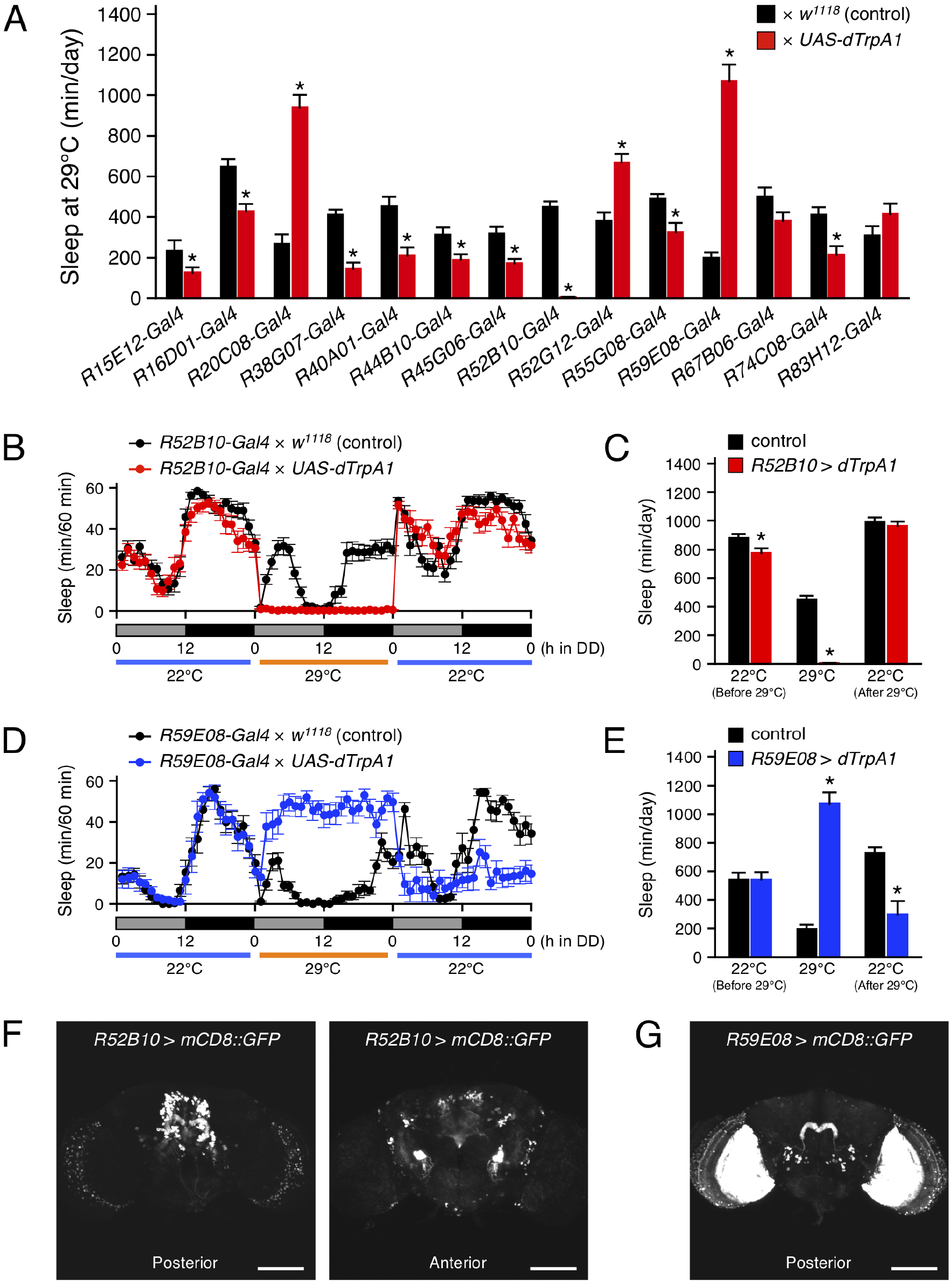
Acute activation of the PB neurons affected sleep. **(A)** The amount of sleep per day at 29°C in DD conditions in control flies (black bars) and flies expressing *dTrpA1* transgene by indicated *Gal4* drivers that highly express in PB neurons (red bars). For controls, each *Gal4* driver was crossed with *w^1118^* (the genetic background of *UAS-dTrpA1* carrying flies). Data are presented as mean ± SEM (*n* = 6-16 for each group). Asterisks indicate statistically significant differences from control determined by a *t*-test (*p* < 0.05). **(B,C)** Sleep profiles for 60-min intervals (B) or total daily sleep (C) for controls (*R52B10-Gal4 × w^1118^*, black circles or bars, *n* = 16) or flies expressing *dTrpA1* with *R52B10-Gal4 (R52B10-Gal4 × UAS-dTrpA1*, red circles or bars, *n* = 16) in DD. Behavior was monitored in DD for 1 day at 22°C, followed by 1 day at 29°C (dTrpA1 activation), and 1 day at 22°C. Gray and black bars under the horizontal axis indicate subjective day and night, respectively. Data are presented as mean ± SEM. \**p* < 0.05 vs. control; *t*-test. **(D,E)** Sleep profiles for 60-min intervals (D) or total daily sleep (E) for controls (*R59E08-Gal4 × w^1118^*, black circles or bars, *n* = 16) or flies expressing *dTrpA1* with *R59E08-Gal4 (R59E08-Gal4 × UAS-dTrpA1*, blue circles or bars, *n* = 10) in DD. Data are presented as mean ± SEM. \**p* < 0.05 vs. control; *t*-test. **(F,G)** Maximum-intensity projection of the confocal brain images of *R52B10-Gal4* (F) or *R59E08-Gal4* (G) crossed to *UAS-mCD8::GFP* flies. The scale bars represent 100 μm.

The expression patterns of the *R52B10-Gal4* or the *R59E08-Gal4* drivers in the adult brains were visualized using *UAS-mCD8::GFP*. The wake-promoting *R52B10-Gal4* drove strong expression in several cell types of columnar neurons named PFN projecting from the PB to the ventral part of the FB and contralateral NO (*Figure 2F*, left panel) as shown in the FlyLight database (Jenett et al., 2012) and the previous report (Buchanan et al., 2015). In addition to these central complex substructures, some neurons in the anterior ventrolateral protocerebrum (AVLP) were also be seen as intense GFP signals (*Figure 2F*, right panel). The thermogenetic activation using the *R52H12-Gal4* or the *R83A10-Gal4* that express strongly in the AVLP neuropil (the FlyLight database, Jenett et al., 2012) did not result in a marked decrease in the amount of sleep (*Figure 2-figure supplement 1*). The sleep-promoting *R59E08-Gal4* specifically labeled the PB strongly in the central complex (*Figure 2G*). These PB neurons were identified as PB interneurons according to their previously described morphological features (Lin et al., 2013; Wolff et al., 2015). In addition, the ubiquitous strong expression of GFP in the medulla cells was also detected in the *R59E08-Gal4* driver. Neuronal activation using two *Gal4* drivers (*R14F09-Gal4* and *R59A12-Gal4*), whose expression patterns in the medulla resemble that of the *R59E08-Gal4* (the FlyLight database, Jenett et al., 2012), did not significantly increase the amount of sleep (*Figure 2-figure supplement 2*).

**Figure 2-figure supplement 1.**
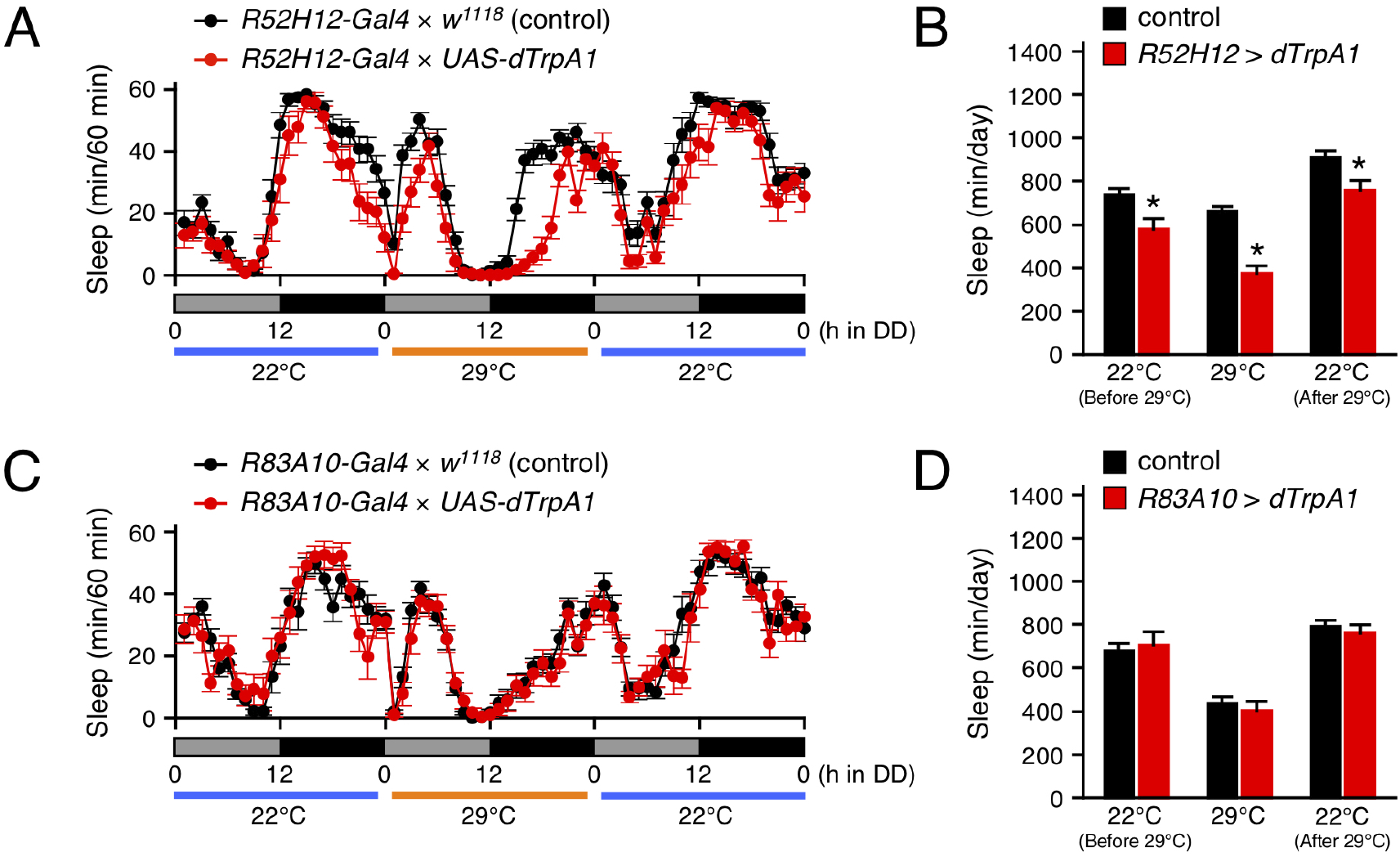
Neuronal activation using *Gal4* drivers that express in the AVLP did not remarkably decrease sleep. **(A,B)** Sleep profiles for 60-min intervals (A) or total daily sleep (B) for controls (*R52H12-Gal4 × w^1118^*, black circles or bars, *n* = 16) or flies expressing *dTrpA1* with *R52H12-Gal4* (*R52H12-Gal4 × UAS-dTrpA1*, red circles or bars, *n* = 14) in DD. The behavior was monitored as described in Figure 2. Data are presented as mean ± SEM. \**p* < 0.05 vs. control; *t*-test. **(C,D)** Sleep profiles for 60-min intervals (C) or total daily sleep (D) for controls (*R83A10-Gal4 × w^1118^*, black circles or bars, *n* = 16) or flies expressing *dTrpA1* with *R83A10-Gal4 (R83A10-Gal4 × UAS-dTrpA1*, red circles or bars, *n* = 11) in DD. Data are presented as mean ± SEM.

**Figure 2-figure supplement 2.**
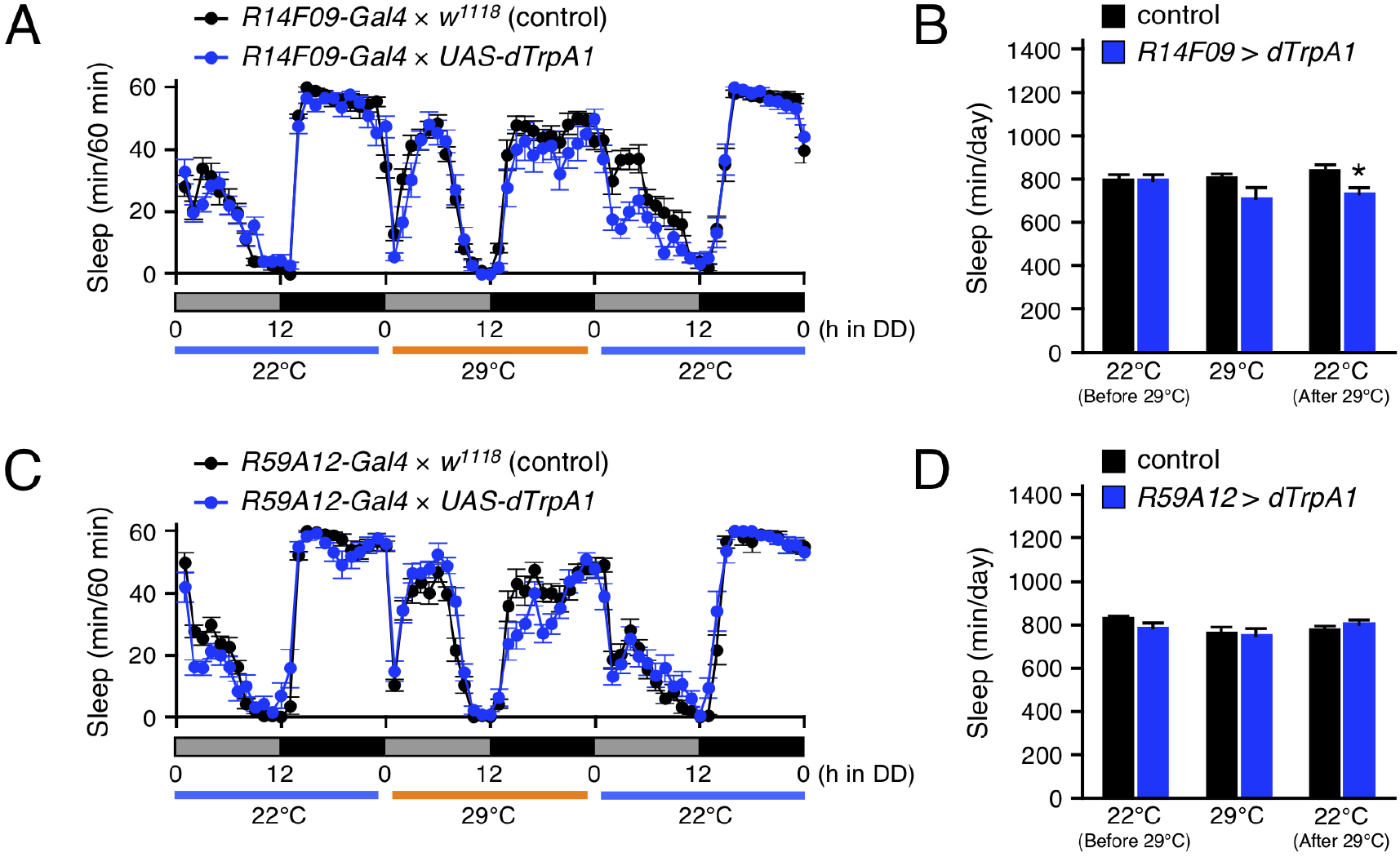
Neuronal activation using *Gal4* drivers that express in the medulla cells did not increase sleep. **(A,B)** Sleep profiles for 60-min intervals (A) or total daily sleep (B) for controls (*R14F09-Gal4 × w^1118^*, black circles or bars, *n* = 16) or flies expressing *dTrpA1* with *R14F09-Gal4* (*R14F09-Gal4 × UAS-dTrpA1*, blue circles or bars, *n* = 16) in DD. The behavior was monitored as described in Figure 2. Data are presented as mean ± SEM. \**p* < 0.05 vs. control; *t*-test. **(C,D)** Sleep profiles for 60-min intervals (C) or total daily sleep (D) for controls (*R59A12-Gal4 × w^1118^*, black circles or bars, *n* = 15) or flies expressing *dTrpA1* with *R59A12-Gal4 (R59A12-Gal4 × UAS-dTrpA1*, blue circles or bars, *n* = 16) in DD. Data are presented as mean ± SEM.

### Activation of PB interneurons with *R59E08-Gal4* increased sleep

The sleep-promoting *R59E08-Gal4* was expressed not only in the adult brain but also in the ventral nerve cord (VNC) (*Figure 3A*). To determine which targeted neurons in this driver are responsible for the sleep induction, we employed *tsh-Gal80*, which inhibits Gal4-mediated transcription in the VNC. The incorporation of the *tsh-Gal80* transgene successfully blocked *R59E08-Gal4* induced GFP expression in the VNC without affecting its expression pattern in the brain (*Figure 3A*). Activation of *R59E08-Gal4*-expressing neurons with *tsh-Gal80* also markedly increased in the amount of sleep, indicating that brain neurons labeled by *R59E08-Gal4* contribute dTrpA1-mediated sleep induction (*Figure 3B,C* and *Figure 2D,E*). On the other hand, decreased sleep caused after dTrpA1 activation was completely restored by introducing *tsh-Gal80*, indicating that *R59E08-Gal4*-expressing the VNC neurons are involved in this sleep phenotype. Thus, distinct populations of *R59E08-Gal4*-expressing neurons participate in the sleep phenotypes induced during or after dTrpA1 activation.

**Figure 3.**
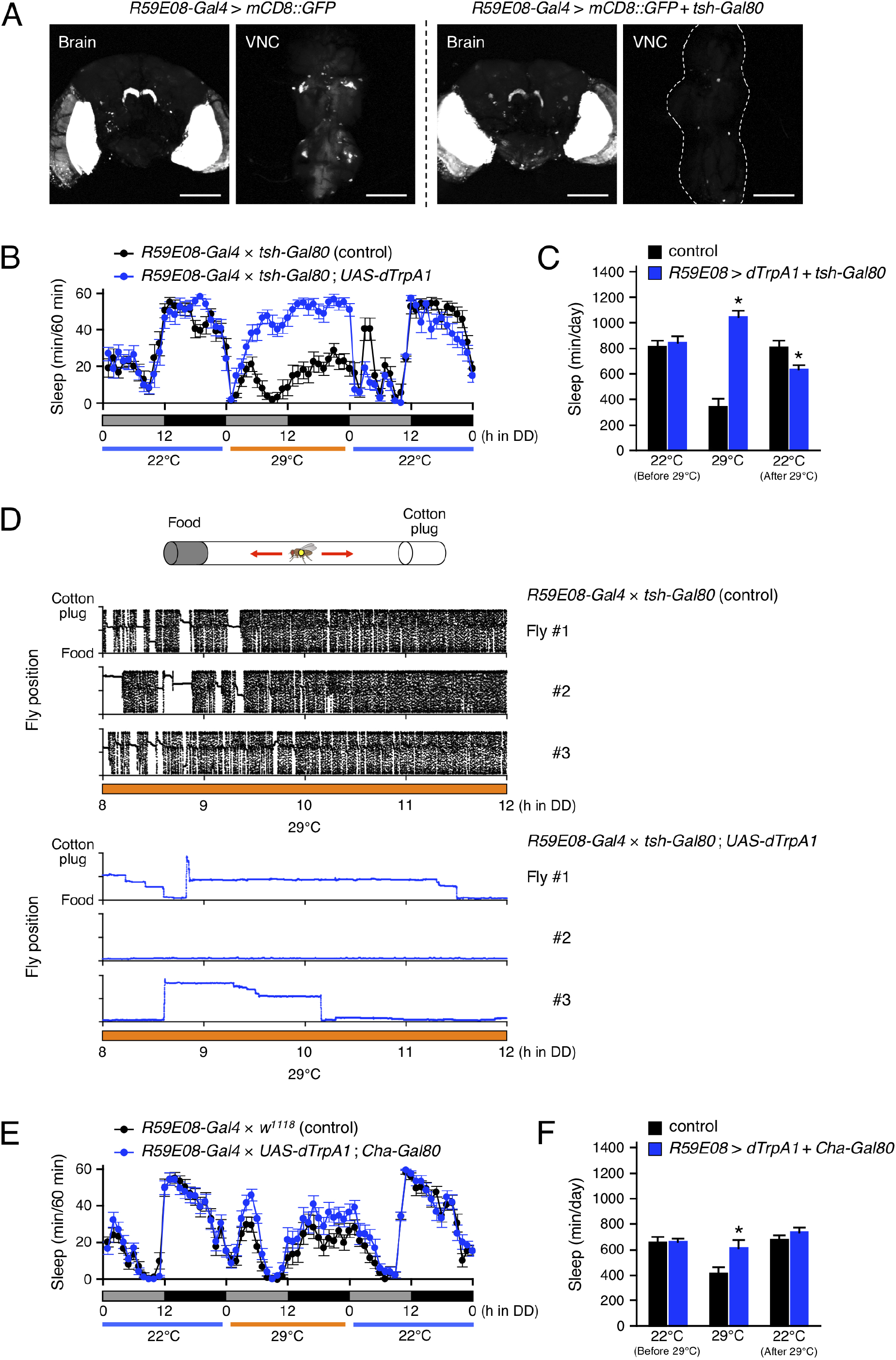
*R59E08-Gal4*-expressing PB interneurons promote sleep. **(A)** Maximum-intensity projection of the confocal brain or ventral nerve cord (VNC) images of flies expressing *UAS-mCD8::GFP* under the control of *R59E08-Gal4* (left two panels) or *R59E08-Gal4* with *tsh-Gal80* (right two panels). Transgene *tsh*-*Gal80* efficiently suppressed *R59E08-Gal4* driven GFP expression in the VNC, while did not in the brains. The scale bars indicate 100 μm. **(B,C)** Sleep profiles in 60-min intervals (B) or total daily sleep (C) for control flies (*R59E08-Gal4 × tsh-Gal80*, black circles or bars, *n* = 12) or flies expressing *dTrpA1* in *R59E08-Gal4* brain neurons (*R59E08-Gal4 × tsh-Gal80*; *UAS-dTrpA1*, blue circles or bars, *n* = 11) in DD. The behavior was monitored as described in Figure 2. Data are presented as mean ± SEM. \**p* < 0.05 vs. control; *t*-test. **(D)** Horizontal movements of the centroid of three representative control flies (*R59E08-Gal4 × tsh-Gal80*, black circles) or flies expressing *dTrpA1* in *R59E08-Gal4* brain neurons (*R59E08-Gal4* × *tsh-Gal80; UAS-dTrpA1*, blue circles) in a glass tube at 29°C during the late subjective day (CT 8 to CT 12). Positions of the cotton plug and the food were at the top and the bottom in each plot, respectively. The behavior of flies was recorded at 2 frames/sec using an infrared video camera in DD conditions. The fly position in a glass tube was calculated for each image using an ImageJ plugin. **(E,F)** Sleep profiles in 60-min intervals (E) or total daily sleep (F) for control flies (*R59E08-Gal4 × w^1118^*, black circles or bars, *n* = 16) or flies expressing *dTrpA1* in *R59E08-Gal4* except cholinergic neurons (*R59E08-Gal4 × tsh-Gal80*; *UAS-dTrpA1*; *Cha^7.4kb^-Gal80*, blue circles or bars, *n* = 16) in DD. Data are presented as mean ± SEM. \**p* < 0.05 vs. control; *t*-test.

Because the DAM system only detects motions of a fly passing through an infrared beam that bisects a glass tube at the center, continuing feeding or grooming away from the beam path are detected as apparent inactivity (continuous bins with the value of zero). To confirm that the flies certainly sleep by activation of *R59E08-Gal4*-expressing brain neurons, we performed video analysis of the behavior of these flies in glass tubes during the late subjective-day (CT 8 to CT 12) at 29°C (*Figure 3D*). The temperature was shifted from 22°C to 29°C at CT 0. As expected from the sleep patterns observed by the DAM system, the experimental flies (*R59E08-Gal4 × tsh-Gal80*; *UAS-dTrpA1*) displayed little locomotor activity compared to controls (*R59E08-Gal4 × tsh-Gal80*). The immobile flies did not show excessive feeding or grooming, and paralysis in naked-eye observation.

We also found that the sleep phenotypes induced during or after activation of *R59E08-Gal4*-expressing neurons were largely rescued by introducing *Cha^7.4kb^-Gal80* transgene, which prohibits Gal4 activity in cholinergic neurons (*Figure 3E,F*). Taken together, these results indicate that the PB interneurons labeled by *R59E08-Gal4* were sleep-promoting and were primarily cholinergic.

### PFN neurons were wake-promoting neurons

As shown in *Figure 4A, tsh-Gal80* effectively attenuated the wake-promoting *R52B10-Gal4* driven GFP expression in the VNC without effect on its expression in the brain. Activation of the brain neurons targeted by *R52B10-Gal4* resulted in a dramatic reduction in the amount of sleep comparable to that evoked by activating both the brain and the VNC neurons using this *Gal4* driver (*Figure 4B,C* and *Figure 2B,C). Cha^7.4kb^-Gal80* was able to fully revert the decreased sleep caused by activation of *R52B10-Gal4-expressing* neurons (*Figure 4D,E* and *Figure 2B,C*). These results suggest that cholinergic brain neurons labeled by *R52B10-Gal4* are wake-promoting.

**Figure 4.**
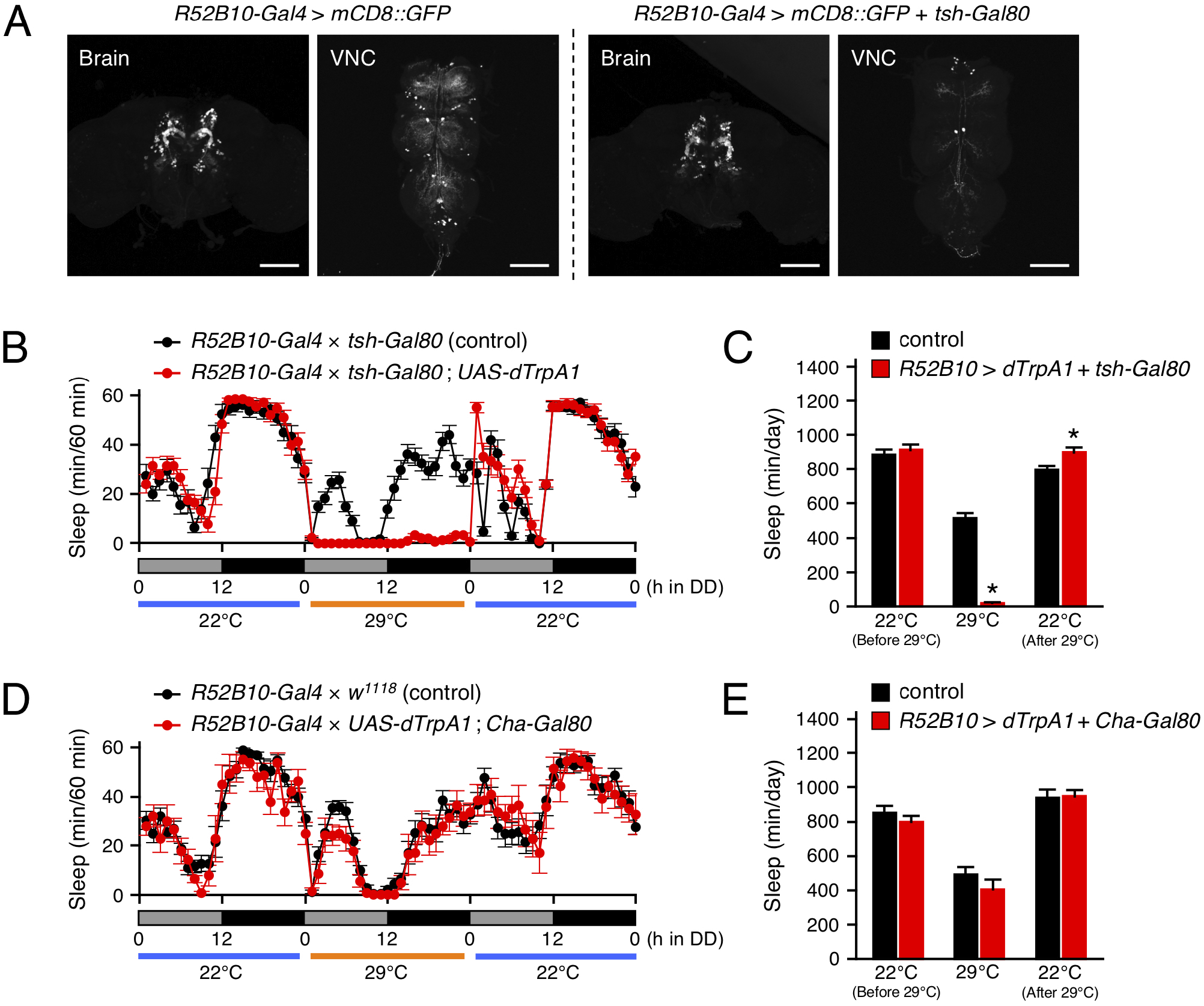
Activation of *R52B10-Gal4*-expressing cholinergic neurons in the brain promotes wakefulness. **(A)** Maximum-intensity projection of the confocal brain or VNC images of flies expressing *UAS-mCD8::GFP* under the control of *R52B10-Gal4* (left two panels) or *R52B10-Gal4* with *tsh-Gal80* (right two panels). The *tsh-Gal80* efficiently suppressed *R52B10-Gal4* driven GFP expression in the VNC, while did not in the brains. **(B,C)** Sleep profiles in 60-min intervals (B) or total daily sleep (C) for control flies (*R52B10-Gal4 × tsh-Gal80*, black circles or bars, *n* = 16) or flies expressing *dTrpA1* in *R52B10-Gal4* brain neurons (*R52B10-Gal4 × tsh-Gal80*; *UAS-dTrpA1*, red circles or bars, *n* = 16) in DD. The behavior was monitored as described in Figure 2. Data are presented as mean ± SEM. \**p* < 0.05 vs. control; *t*-test. **(D,E)** Sleep profiles in 60-min intervals (D) or total daily sleep (E) for control flies (*R52B10-Gal4 × w^1118^*, black circles or bars, *n* = 16) or flies expressing *dTrpA1* in *R52B10-Gal4* except cholinergic neurons (*R52B10-Gal4 × UAS-dTrpA1*; *Cha^7.4kb^-Gal80*, red circles or bars, *n* = 7) in DD. Data are presented as mean ± SEM.

Next, we searched for *R52B10-Gal4*-expressing brain neurons that contribute to the promotion of wakefulness by a genetic mosaic approach. Using the mosaic analysis with a repressible cell marker (MARCM) system, the *dTrpA1* and *GFP* were randomly co-expressed in a subpopulation of neurons using *R52B10-Gal4* with FRT/FLP-induced mitotic recombination. We measured the amount of sleep in these genetically mosaic flies at 22°C and then 29°C under DD conditions. The change in sleep amount (ΔSleep) in individual flies was calculated by subtracting the amount of sleep during the 12-h subjective night at 29°C from that at 22°C (*Figure 5A*). In this experiment, ΔSleep value above the mean plus two times standard deviation (+2 SD) was regarded as a significant decrease in subjective-night sleep at 29°C. Of 133 flies examined, 7 flies had a ΔSleep value above the mean +2 SD (426 min). After measuring the amount of sleep, the brains of these flies were dissected and immunostained to determine GFP expression co-expressed with dTrpA1. GFP expression in the flies with significantly reduced sleep labeled the PFN neurons in all but one of the flies shown in #7 in *Figure 5B*. By contrast, in the 7 individuals having ΔSleep value near the average, 2 flies were labeled with the PFN neurons (#5 and #6 in *Figure 5C*), while the remaining flies were either not labeled at all (#1 and #2) or labeled with neurons different from the PFN neurons (#3, #4, #7).

**Figure 5.**
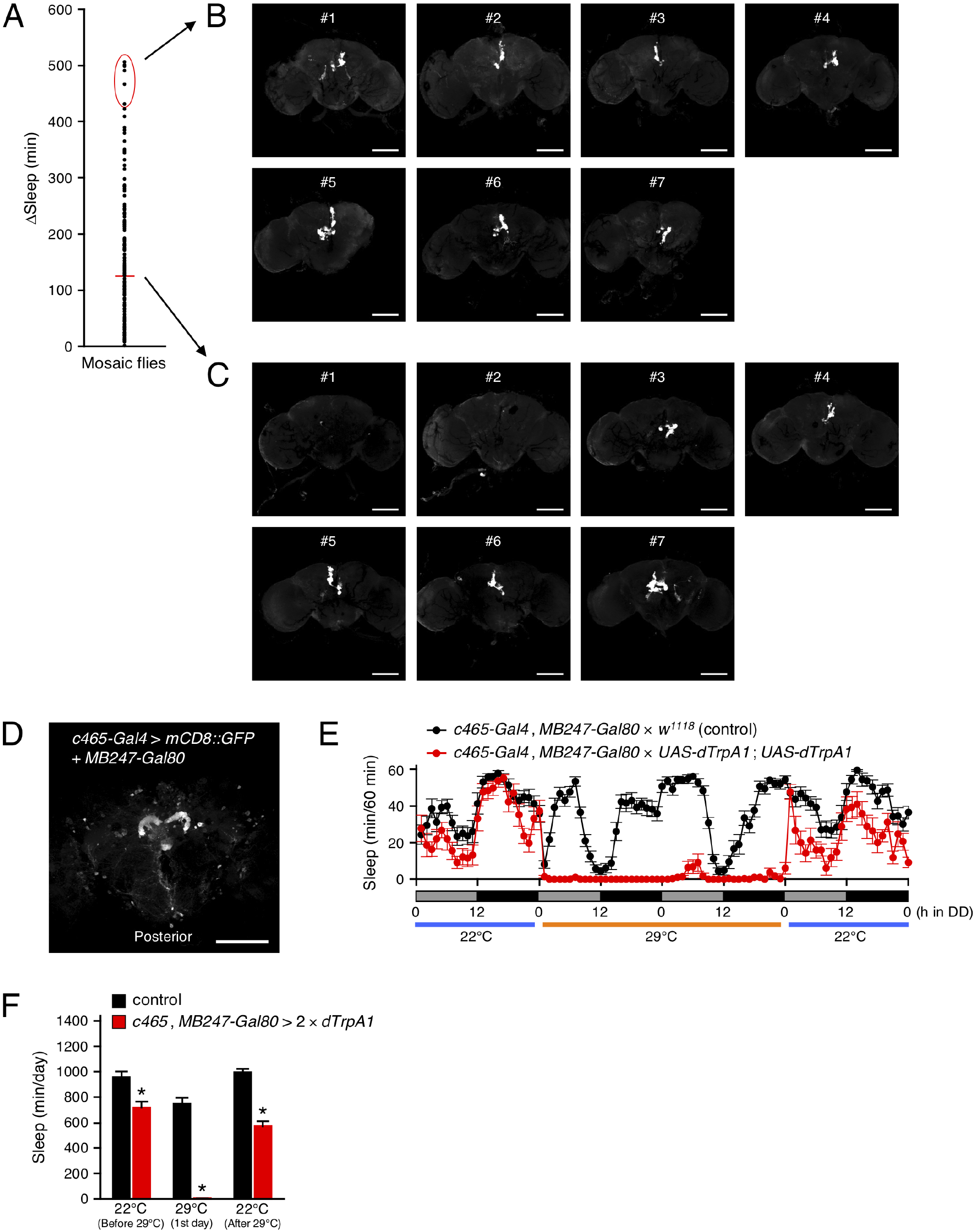
*R52B10-Gal4*-expressing PFN neurons promote wakefulness. **(A)** Using the MARCM system, *dTrpA1* expression was targeted to a limited number of neurons in *R52B10-Gal4*. The behavior of mosaic flies was monitored as described in Figure 2. Sleep change (ΔSleep) of a single fly was calculated by subtracting the amount of sleep during the subjective night at 29°C from that at 22°C (before 29°C) (*n* = 133). The red horizontal bar indicates the mean. **(B,C)** Maximum-intensity projection of the confocal brain images of the flies whose ΔSleep were > +2 SD higher than the mean (red oval in (A)) (B) or nearly equal to the mean (C). **(D)** Maximum-intensity projection of the confocal brain images of *c465-Gal4, MB247-Gal80* crossed to *UAS-mCD8::GFP* flies. *c465-Gal4* drives expression in the PFN neurons. **(E,F)** Sleep profiles in 60-min intervals (E) or total daily sleep (F) for control (*c465-Gal4*, *MB247-Gal80 × w^1118^*, black circles or bars, *n* = 15) or flies expressing *dTrpA1* in *c465-Gal4* except for mushroom body neurons (*c465-Gal4, MB247-Gal80 × UAS-dTrpA1*; *UAS-dTrpA1*, red circles or bars, *n* = 11) in DD conditions. Behavior was monitored as described in Figure 2, except that flies were transferred to 29°C for 2 days to allow dTrpA1 activation. Data are presented as mean ± SEM. \**p* < 0.05 vs. control; *t*-test. All of the scale bars indicate 100 μm.

Moreover, to ensure that the PFN neurons labeled by *R52B10-Gal4* are the wake-promoting neurons, we employed *c465-Gal4* that drives expression in these neurons participating in the tuning of the magnitude of locomotor handedness (Buchanan et al., 2015). Because *c465-Gal4* is also strongly expressed in the mushroom bodies that is another brain neuropil structures involved in *Drosophila* sleep regulation (Joiner et al., 2006; Pitman et al., 2006; Sitaraman et al., 2015a), we used *MB247-Gal80*, which blocks the activity of Gal4 in the mushroom bodies (*Figure 5D*). As expected, activation of *c465-Gal4*-expressing neurons except for mushroom body neurons significantly decreased sleep, as did the activation of *R52B10-Gal4*-expressing neurons (*Figure 5E,F*). These results indicate that the PFN neurons labeled by *R52B10-Gal4* are wake-promoting.

### The sleep-promoting PB interneurons had synaptic connections with the wake-promoting PFN neurons

To visualize putative dendritic and axonal terminals of the wake-promoting PFN neurons, both DenMark (Nicolai et al., 2010), a specific somatodendritic marker and synaptotagmin (syt)-GFP (Zhang et al., 2002), a presynaptic marker localized to synaptic vesicles were expressed using *R52B10-Gal4*. These neurons displayed specific dendritic DenMark labeling in the PB and strong labeling of presynaptic syt-GFP in the layer 1 of the FB and in the ventral NO (*Figure 6A*). In addition, weak labeling of presynaptic terminals was also observed near the layers 2 to 3 of the FB and in regions of the NO other than the ventral NO.

**Figure 6.**
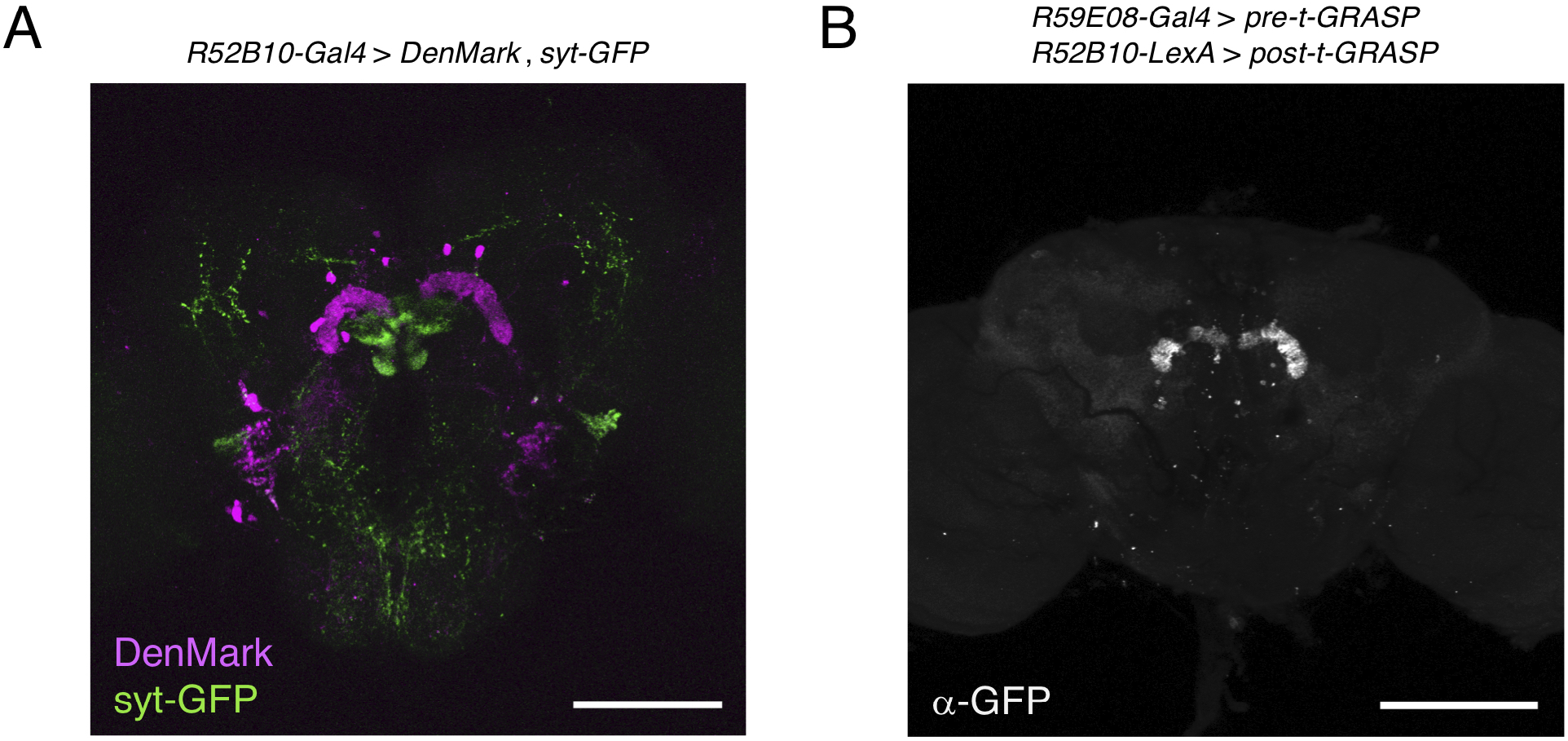
The sleep-promoting PB interneurons form synaptic contacts with the wake-promoting PFN neurons. **(A)** The dendritic arbors and presynaptic terminals of the PFN neurons in *R52B10-Gal4* were visualized by expression of the postsynaptic marker DenMark and the presynaptic marker syt-GFP, respectively. **(B)** The neuronal connection of the sleep-promoting PB interneurons to the wake-promoting PFN neurons was revealed using the t-GRASP method. The *pre-t-GRASP* and the *post-t-GRASP* encode the split-GFP fragments, which are targeted to presynaptic endings and dendritic terminals, respectively. Brains expressing *pre-t-GRASP* with *R59E08-Gal4* and *post-t-GRASP* with *R52B10-LexA* were stained with a reconstituted GFP-specific antibody. The scale bars represent 100 μm.

To assess synaptic connectivity between the sleep-promoting PB interneurons and the wake-promoting PFN neurons, we employed a targeted GFP reconstitution across synaptic partners (t-GRASP) technique (Shearin et al., 2018) under the control of two independent binary systems. A part of the *cacophony* gene tagged with the *GFP11* fragment of a *split-GFP* (*pre-t-GRASP*), which is targeted to axonal terminals and a portion of the mouse *Icam5* gene tagged with the *GFP1-10* (*post-t-GRASP*), which is targeted to dendritic terminals were expressed by *R59E08-Gal4* and *R52B10-LexA*, respectively. Reconstituted GFP signals were detected in the PB (*Figure 6B*), suggesting that the sleep-promoting PB interneurons labeled by *R59E08-Gal4* have synaptic connections with the wake-promoting PFN neurons marked by *R52B10-Gal4*.

### Dopamine signaling acted on the sleep-promoting PB interneurons for sleep regulation

Dopamine has been identified as a key neurotransmitter in the regulation of sleep in *Drosophila* (Andretic et al., 2005; Kume et al., 2005; Pimentel et al., 2016). In the *Drosophila* brain, DA neurons are distributed in clusters, and each cluster is involved in various physiological phenomena, including sleep-wake (Liu et al., 2012; Ueno et al., 2012; Sitaraman et al., 2015b). A pair of DA neurons, named T1, has been previously observed with dendrites arborizing in the tritocerebrum and with axons projecting to the PB (Nässel and Elekes, 1992; Alekseyenko et al., 2013). The intersectional method combining the enhancer-trap flippase (FLP) transgenic line, *FLP^243^* with a DA-neuron specific *TH-Gal4* driver allows for the selective targeting of the T1 neurons (*Figure 7A*) (Alekseyenko et al., 2013). Both activation and inhibition of the T1 DA neurons using this intersectional method have been reported to specifically facilitate inter-male aggression. On the other hand, inactivation of these neurons has no effect on locomotor activity and sleep amount (Alekseyenko et al., 2013). We conditionally activated the T1 neurons with dTrpA1 and found that the manipulation resulted in a significant decrease in the amount of sleep (*Figure 7B,C*).

**Figure 7.**
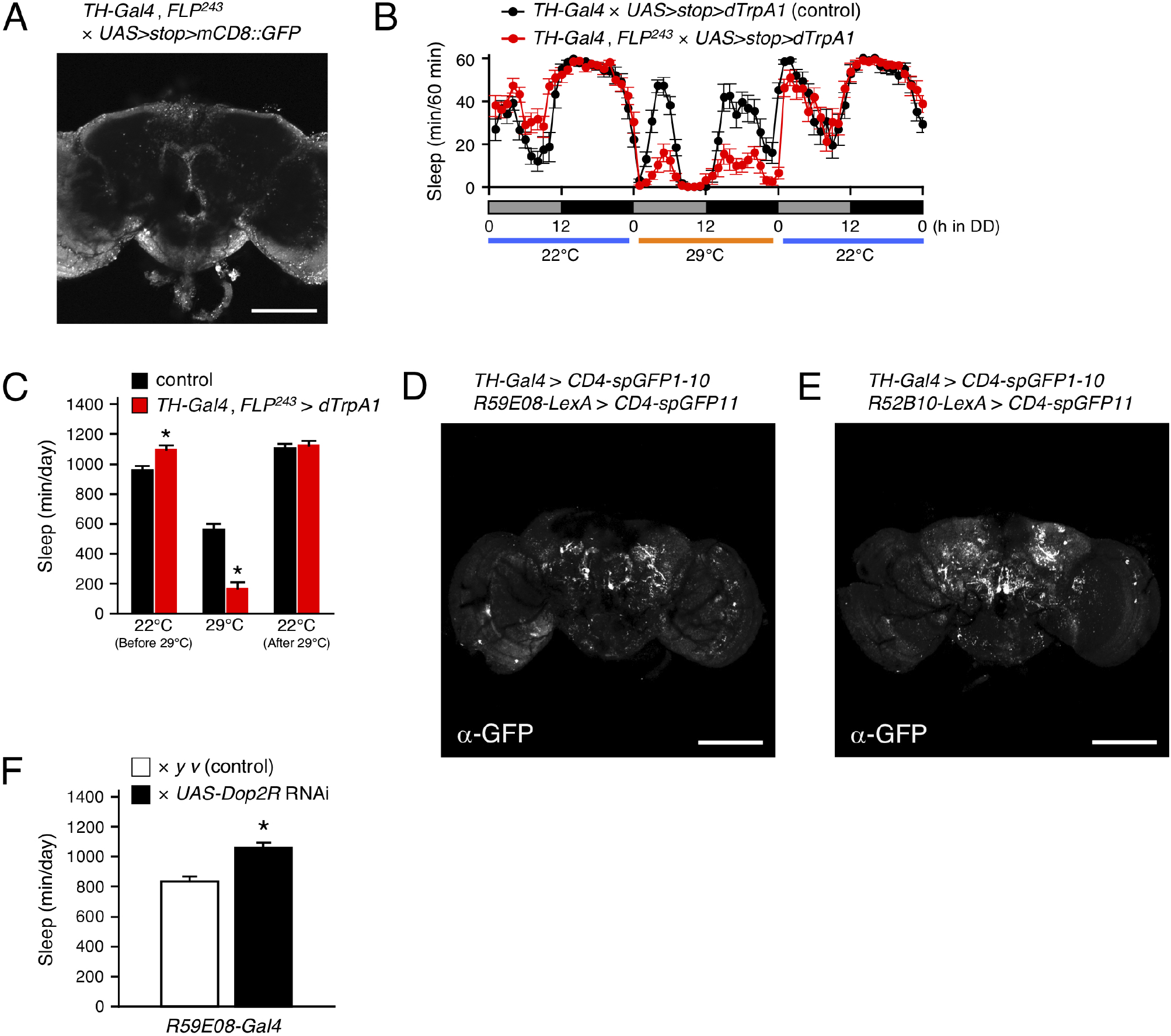
Dopamine regulates sleep by acting on the sleep-promoting PB interneurons. **(A)** Maximum-intensity projection of the confocal brain images of *TH-Gal4, FLP^243^* crossed to *UAS>stop>mCD8::GFP* flies. *TH-Gal4, FLP^243^* restricts expression to the T1 dopaminergic (DA) neurons that project to the PB. **(B,C)** Sleep profiles in 60-min intervals (B) or total daily sleep (C) for control (*TH-Gal4 × UAS>stop>dTrpA1*, black circles or bars, *n* = 15) or flies expressing *dTrpA1* in T1 DA neurons (*TH-Gal4, FLP^243^ × UAS>stop>dTrpA1*, red circles or bars, *n* = 16) in DD conditions. The behavior was monitored as described in Figure 2. Data are presented as mean ± SEM. \**p* < 0.05 vs. control; *t*-test. **(D,E)** The anatomical connection of DA neurons in *TH-Gal4* to the sleep-promoting PB interneurons (D) or the wake-promoting PFN neurons (E) were examined using the GRASP method. Brains were stained with a reconstituted GFP-specific antibody. **(F)** Total daily sleep for control (white bar, *n* = 15) and *Dop2R* RNAi-expressing flies using the *R59E08-Gal4* driver (black bars, *n* = 16) in DD conditions. Data are averaged for 3 days and are presented as mean ± SEM. \**p* < 0.05; *t*-test. All of the scale bars indicate 100 μm.

Next, we examined whether T1 DA neurons directly connect with sleep-wake regulating PB neurons by GRASP analysis utilizing membrane-tethered two complementary split-GFP fragments (CD4-spGFP1-10 and CD4-spGFP11) (Feinberg et al., 2008). *TH-Gal4* was used to express *UAS-CD4-spGFP1-10* and *R59E08-LexA* or *R52B10-lexA* were employed to drive *lexAop-CD4-spGFP11*, respectively. Immunohistochemical signals for reconstituted GFP in the PB were detected only in the combination of *TH-Gal4* and *R59E08-LexA (Figure 7D,E*). These signals were dot-like and localized to the bilateral bends of the PB.

Alekseyenko and colleagues also demonstrate that Dopamine 2-like receptors (Dop2R also known as D2R or DD2R), a functional counterpart of mammalian D2 receptor, are expressed in the PB but not localized to the presynaptic terminals of T1 neurons. *Dop2R* knockdown using two independent RNAi lines in the *R59E08-Gal4* expressing PB neurons resulted in a significant increase in the amount of sleep compared to controls (*Figure 7F* and *Figure 7-figure supplement 1*). These results support the idea that T1 DA neurons regulate sleep by acting on the sleep-promoting PB interneurons via D2 receptor signaling.

**Figure 7-figure supplement 1.**
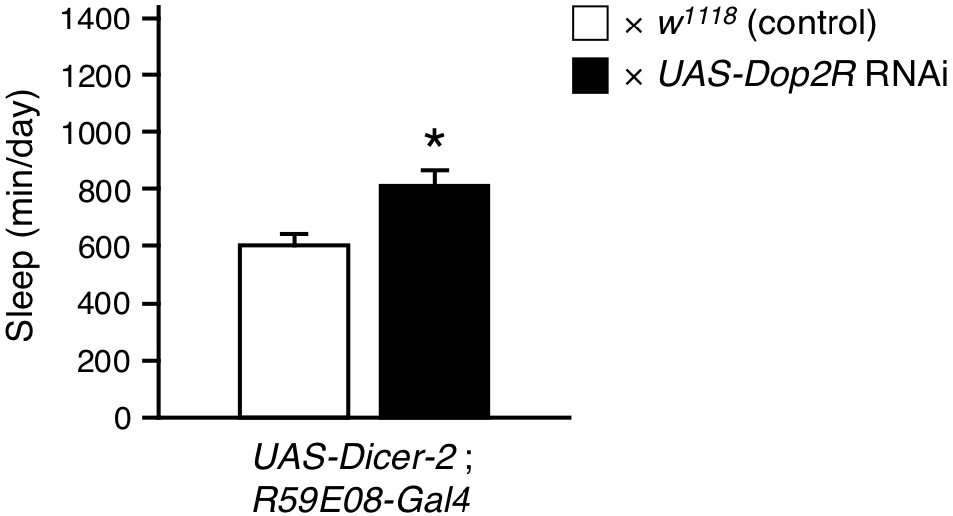
*Dop2R* knockdown with *R59E08-Gal4* increased sleep. Total daily sleep for control (*UAS*-*Dicer*-*2*; *R59E08-Gal4 × w^1118^*, white bar, *n* = 13) and *Dop2R* RNAi (VDRC 1820)-expressing flies using the *R59E08-Gal4* driver (*UAS*-*Dicer*-*2*; *R59E08-Gal4 × UAS-Dop2R* RNAi, black bars, *n* = 13) in DD conditions. Data are averaged for 3 days and are presented as mean ± SEM. \**p* < 0.05; *t*-test.

## Discussion

The data presented provide evidence that *Drosophila* sleep is controlled by the neuronal circuit in PB consisting of the wake-promoting PFN neurons and the sleep-promoting PB interneurons and by the T1-PB dopamine pathway (*Figure 7-figure supplement 2*). Previous anatomical, electrophysiological and genetic studies have revealed the physiological functions of the PB in several insect species. For example, in desert locusts, different types of PB neurons generate a topographical map of the sky polarization pattern underlying sun compass orientation (Homberg et al., 2011). Morphological counterparts of these locust’s PB neurons are found in migratory monarch butterflies and serve similar functions in processing polarized-light information (Heinze and Reppert, 2011). Experiments in fruit flies have demonstrated a role for the PB in locomotor control such as walking speed, leg coordination, maintenance of walking activity, locomotor handedness and visual targeting in gap-crossing behavior (Strauss et al., 1992; Strauss and Heisenberg, 1993; Martin et al., 1999; Triphan et al., 2010; Buchanan et al., 2015). The locomotor activity of *nob^KS49^* mutants has been reported to be decreased compared to control flies but was increased in the present study (Martin et al., 1999) (*Figure 1A*). We and Martin et al. assessed the number of activity counts per day or at a particular time of day (4.5 hours) in *nob^KS49^* flies as total locomotor activity, respectively. This difference in assessment methods may have caused the phenotypic discrepancy in locomotor activity. *nob^KS49^* flies with a large gap at the sagittal midplane in the PB showed a significant reduction in the amount of sleep (*Figure 1B*), suggesting a physical impairment of sleep-promoting neurons in this mutant. We found that *R59E08-Gal4*-expressing PB interneurons promote sleep (*Figure 2* and *Figure 3*). Analysis of single PB neurons labeled by MARCM or multicolor flip-out technique has shown that PB interneurons are classified into three cell types based on their pattern of glomerular innervation in the PB (Lin et al., 2013; Wolff et al., 2015). Two of the three cell types of PB interneurons innervate all glomeruli, including the central part of the PB. Because the sleep-promoting *R59E08-Gal4* was expressed throughout the PB, at least one of these two cell types of PB interneurons would have been labeled with this driver. Although the cell type of PB interneurons that promote sleep has been unidentified in our results, the two cell types may be physically damaged in *nob^KS49^* mutants.

Our MARCM experiment using *R52B10-Gal4* suggested that distinct types of PFN neurons promote wakefulness (*Figure 5B*). There are five cell types of PFN neurons, which are PB output neurons, classified according to their patterns of projection to the FB and the NO (Wolff et al., 2015). Each PFN neuron belonging to the same cell type is further classified into subtypes that have different patterns of arborization in the PB and arborize to one of the PB glomeruli. The *Gal4* line *c465* drives expression in the wake-promoting PFN neurons (*Figure 5D-F*) (Lin et al., 2013; Buchanan et al., 2015). Of the five cell types of PFN neurons, three are commonly labeled by *R52B10-Gal4* and *c465-Gal4*, especially the cell type with axon terminals in layer 2 of the FB and in the ventral NO exhibits high *Gal4* expression in these two drivers (Buchanan et al., 2015). On the other hand, neuronal activation with *R44B10-Gal4*, which is highly expressed in all three cell types of PFN neurons (Buchanan et al., 2015), was less effective in reducing the amount of sleep than neuronal activation with *R52B10-Gal4* or *c465-Gal4* (*Figure 2A*). Although *R52B10-Gal4*, *c465-Gal4* and *R44B10-Gal4* label the same cell types of PFN neurons, it is possible that the subtypes of PFN neurons arborizing in one specific PB glomerulus which are co-labeled with *R52B10-Gal4* and *c465-Gal4* are implicated in promoting arousal.

The t-GRASP experiment using *R59E08-Gal4* and *R52B10-LexA* suggested the sleep-promoting PB interneurons form synaptic contacts with the wake-promoting PFN neurons (*Figure 6B*). The symmetrical patterns of the t-GRASP signal intensity in PB glomeruli should indicate that this signal is not non-specific, supporting the idea described above that particular subtypes of PFN neurons promote arousal. Sleep phenotypes of the flies in which *R59E08-Gal4-* or *R52B10-Gal4*-expressing neurons were acutely activated, and axon-dendrite connectivity between these sleep-regulating neurons suggest that the sleep-promoting PB interneurons inhibit the activity of the wake-promoting PFN neurons. Because the sleep-promoting PB interneurons were predominantly cholinergic (*Figure 3E,F*), the wake-promoting PFN neurons were likely to be regulated by inhibitory acetylcholine receptor signaling (Ren et al., 2015). Functional connectivity analysis of the central complex using Ca^2+^ imaging combined with optogenetics has demonstrated that activation of one cell type of PB interneurons designated as Δ7 triggers the inhibitory response of one cell type of PFN neurons (Franconville et al., 2018). However, since the Δ7 neurons are either glutamatergic or GABAergic, they must not be the major PB interneurons to promote sleep. Further anatomical and functional dissection of the sleep-regulating neuronal circuits in the PB archived with promising *split-Gal4* lines will be required.

Previous studies have elucidated the control of *Drosophila* sleep by several clusters of DA neurons (Liu et al., 2012; Ueno et al., 2012; Sitaraman et al., 2015b). This study revealed that the PB-projecting T1 DA neurons physically connected to the sleep-promoting PB interneurons promote wakefulness (*Figure 7A-D*). Because knockdown of *Dop2R* encoding Gi protein-coupled dopamine receptor in *R59E08-Gal4*-expressing neurons significantly increased the amount of sleep (*Figure 7F* and *Figure 7-figure supplement 1*), the simplest explanation is that dopamine inhibits the sleep-promoting PB interneurons and thus promotes wakefulness. *Dop2R* null mutants have an increased amount of sleep (Deng et al., 2019). The DopR2 function in the pars intercerebralis (PI) neurons expressing Dilp2 and SIFamide has been shown to contribute to this sleep-increasing phenotype. Our results show that, in addition to the PI, Dop2R also functions in the PB to regulate sleep.

T1 neurons were originally identified as neurons that specifically modulate aggression between pairs of males (Alekseyenko et al., 2013). Regarding the interaction between sleep and aggression, it has been shown that aggressive behaviors are suppressed in sleep-deprived male flies, and the changes are mediated by octopamine signaling (Kayser et al., 2015). Interestingly, Duhart and colleagues have recently reported that T1 neurons act downstream of courtship- and sleep-regulating P1 neurons to modulate nutrition-dependent sleep-courtship balance in male flies (Duhart et al., 2020). Elucidation of the mechanisms controlling the activity of T1 neurons may provide a novel link between sleep, aggression and courtship.

More recently, we have successfully demonstrated that the wake-promoting PFN neurons directly activate the FB interneurons (also known as pontine neurons) that appear to transmit inhibitory acetylcholine signals to the sleep-promoting dFB neurons (Kato YS et al., unpublished results, *Figure 7-figure supplement 2*). Further studies are required to determine the cellular and molecular details of how these PB-dFB circuits control sleep-wakefulness.

**Figure 7-figure supplement 2.**
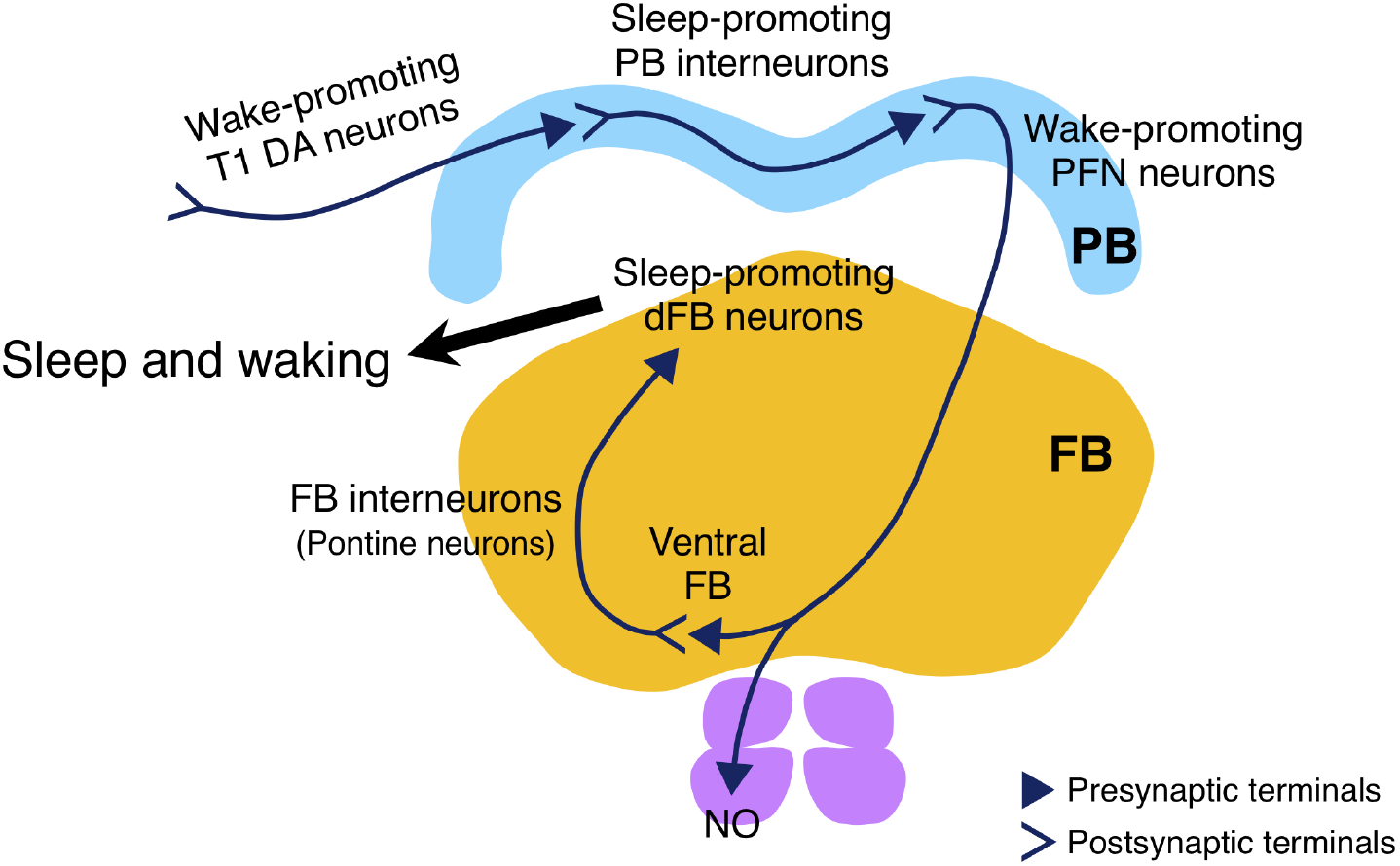
Schematic drawing of the sleep-regulating neuronal circuit in the PB. The circuit consists of the sleep-promoting PB interneurons and the wake-promoting PFN neurons, and is regulated by the wake-promoting T1 DA neurons. The wake-promoting PFN neurons indirectly inhibit the sleep-promoting dFB neurons via the FB interneurons (Kato YS et al., unpublished results).

## Supporting information

PB-sleep_Fig2_supplement1

PB-sleep_Fig2_supplement2

PB-sleep_Fig7_supplement1

PB-sleep_Fig7_supplement2

## Acknowledgments

We thank Drs. Roland Strauss, Paul A. Garrity, Julie H. Simpson, Takaomi Sakai, J. Douglas Armstrong, Edward A. Kravitz and Barry J. Dickson, the Bloomington *Drosophila* Stock Center, the Vienna *Drosophila* RNAi Center and the Kyoto Stock Center for providing fly lines used in this study. In addition, we also thank Dr. Taro Ueno (Toho University) and members of the Kume lab for helpful discussions. This work was supported by Japan Society for the Promotion of Science (JSPS) KAKENHI Grant Numbers JP18H02481, K.K., JP26507008, JP17K08571, J.T.

## Competing interests

The authors declare that no competing interests exist.

## Materials and Methods

### Fly strains and rearing conditions

Fruit flies (*Drosophila melanogaster*) were raised at 25°C in 50-60% relative humidity on standard medium containing cornmeal, yeast, glucose, wheat germ and agar. They were maintained under a 12-h light: 12-h dark (LD) cycle. Eleven mutant flies with structural defects in the central complex *agnostic^X1^* (*agn^X1^*), *central-body-defect^KS171^* (*cbd^KS171^*), *cbd^KS188^, cbd*^762^, *central-complex-broad^KS145^* (*ccb^KS145^*), *central-complex-deranged^KS135^* (*ccd^KS135^*), *central-complex^KS181^* (*cex^KS181^*), *ellipsoid-body-open^KS263^* (*ebo^KS263^*), *ebo^678^, ebo^1041^* and *no-bridge^KS49^* (*nob^KS49^*) were kindly provided by Roland Strauss. *UAS-dTrpA1* lines on the second or third chromosomes were gifts from Paul A. Garrity and backcrossed over five generations to the control strain (*w^1118^*). *teashirt* (*tsh*)*-Gal80/CyO* (Clyne and Miesenböck, 2008) was a gift from Julie H. Simpson. *Cha^7.4kb^-Gal80* (Sakai et al., 2009) and *MB247-Gal80* (Krashes et al., 2007) were gifts from Takaomi Sakai. *c465-Gal4* (Young and Armstrong, 2010) was a gift from J. Douglas Armstrong. *FLP^243^* (Alekseyenko et al., 2013) was a gift from Edward A. Kravitz. *UAS>stop>mCD8::GFP* (Yu et al., 2010) and *UAS>stop>dTrpA1* (von Philipsborn et al., 2011) were gifts from Barry J. Dickson. *R14F09-Gal4* (stock number: 48652), *R15E12-Gal4* (#48608), *R16D01-Gal4* (#48722), *R37G05-Gal4* (#48133), *R38G07-Gal4* (#50019), *R40A01-Gal4* (#50072), *R44B10-Gal4* (#50202), *R45G06-Gal4* (#50244), *R52B10-Gal4* (#38820), *R52G12-Gal4* (#49581), *R52H12-Gal4* (#38856), *R55G08-Gal4* (#50422), *R59A12-Gal4* (#39206), *R59E08-Gal4* (#39219), *R67B06-Gal4* (#48294), *R74C08-Gal4* (#46711) *R83A10-Gal4* (#48371), *R83H12-Gal4* (#40374), *R91A12-Gal4* (#40573), *10×UAS-IVS-mCD8::GFP* (#32187), *10×UAS-IVS-mCD8::GFP* (#32188), *UAS-DenMark, UAS-syt.eGFP* (#33064), *R52B10-LexA* (#52826), *R59E08-LexA* (#52832), *13×LexAop2-post-t-GRASP, 20×UAS-pre-t-GRASP* (#79040), *UAS-CD4-spGFP1-10, lexAop-CD4-spGFP11* (#58755), *UAS-Dop2R-RNAi* (#50621) and *y*, *v; P{CaryP}attP40* (control line for TRiP RNAi lines, #36304) were obtained from the Bloomington *Drosophila* Stock Center, Indiana University, Indiana, USA. *UAS-Dop2R-RNAi* (VDRC ID: 1820), *UAS*-*Dicer*-*2* (60008) and *w^1118^* (the genetic background for VDRC RNAi lines, 60000) were from the Vienna *Drosophila* RNAi Center (VDRC). *P{neoFRT}19A* (stock number: 106482) and *P{neoFRT}19A, tubP-GAL80, hsFLP;Pin*^YT^/CyO (#108064) were from the Kyoto Stock Center, Kyoto Institute of Technology, Kyoto, Japan.

### Locomotor activity and sleep analysis

Locomotor activity of individual fly was measured using the *Drosophila* activity monitoring system (TriKinetics, Waltham, MA) as described previously. Flies were placed individually in glass tubes (length, 65 mm; inside diameter, 3 mm) containing either standard medium or sucrose-agar medium (5% sucrose and 1% agar) at one end and were entrained for at least 3 days to LD cycles before transferring to constant dark (DD) conditions. Two-to four-day-old male flies were used except for MARCM experiments. Activity data were recorded continuously at 1-min intervals under both LD and DD conditions. *Drosophila* sleep was defined as continuous immobile periods lasting 5 min or longer based on previous reports (Hendricks et al., 2000; Shaw et al., 2000; Huber et al., 2004; Kume et al., 2005). Total activity counts, total amount of sleep and waking activity index were analyzed by Microsoft Excel-based software as previously described (Kume et al., 2005) and averaged over 3 days. For conditional dTrpA1 activation experiments, control and experimental flies were grown at 22°C. Sleep measurements were performed in DD following LD cycles. At circadian time (CT) 0 (the beginning of a subjective day), temperature was raised from 22°C to 29°C for 24 h or 48 h to activate dTrpA1-expressing neurons and then returned to 22°C.

### Video tracking of locomotion

Offspring from the control (*R59E08-Gal4 × tsh-Gal80*) and the experimental (*R59E08-Gal4 × tsh-Gal80; UAS-dTrpA1*) crosses were housed in glass tubes and entrained to an LD cycle at 22°C. After entrainment, these flies were maintained in DD with illumination by near-infrared LEDs. At CT 0 on day 2 in DD, flies were transferred from 22°C to 29°C for 12 h to activate dTrpA1. The behavior of flies was recorded at 2 frames/sec using a USB camera (Flea3 USB3.0: FL3-U3-13Y3M-C, Point Grey Research Inc., Richmond, BC, Canada) fitted with a macro zoom lens (MLH-10X, computer, Tokyo, Japan) during late subjective day (CT 8 to CT 12) at 29°C. The captured images were converted into 8-bit grayscale images. A composite background image was subtracted from each video image. The position of individual fly was calculated for each subtracted image using the Particle Tracker 2D/3D, an ImageJ plugin (Sbalzarini and Koumoutsakos, 2005).

### The MARCM system

To generate flies for MARCM, female flies carrying *FRT19A, tub-GAL80, hs-FLP; UAS-dTrpA1* (X; II) crossed to males carrying *FRT19A; R52B10-Gal4. 10×UAS-IVS-mCD8::GFP* (X; III) at 22°C. Following egg-laying for 48 h, embryos or first instar larvae were heat-shocked at 37°C for 1 h to induce FLP-mediated recombination across two FRT sites. The amount of sleep in mosaic female flies was measured at 22°C and 29°C under DD conditions. After sleep analysis, GFP expression co-expressed with dTrpA1 in the brains was immunohistochemically determined.

### Immunohistochemistry

To detect GFP expression in the MARCM experiments and reconstituted GFP signals in the t-GRASP and GRASP experiments, whole-mount immunofluorescence staining of adult brains was performed as described (Wu and Luo, 2006). For determination of GFP expression, GFP polyclonal antibody (A6455, Thermo Fisher Scientific) was used at 1:250 dilution and goat anti-rabbit IgG, Alexa Fluor 568 (A11036, Thermo Fisher Scientific) was used at 1:200 dilution as secondary antibodies. We used GFP recombinant rabbit monoclonal antibody (G10362, Thermo Fisher Scientific) at 1:300 dilution and monoclonal anti-GFP, clone GFP-20 (G6539, Sigma-Aldrich) at 1:100 dilution for detection of reconstituted GFP signals in t-GRASP and GRASP experiments, respectively. Alexa Fluor 488, goat anti-rabbit IgG (A11008, Thermo Fisher Scientific) and goat anti-mouse IgG (A11001, Thermo Fisher Scientific) were used at 1:200 dilution as secondary antibodies.

### Confocal imaging

Immunostained brain tissues were imaged by laser scanning confocal microscopes Zeiss LSM 510 or LSM 800 (Carl Zeiss). Adult brains and ventral nerve cords expressing *mCD8::GFP* or *syt-GFP* under the control of Gal4 were dissected and scanned by the confocal microscopes without staining. In Figure 3A and Figure 4A, to examine the inhibitory effect of *tsh-Gal80* on the Gal4 activity of *R59E08-Gal4* and *R52B10-Gal4*, the Gal4-driven *mCD8::GFP* expression in the brains or the ventral nerve cords in the presence and absence of *tsh-Gal80* were imaged and processed in parallel using identical settings on a confocal microscope.

### Statistical analysis

Data were analyzed as described in the figure legends using Microsoft Excel and R: a language and environment for statistical computing (R Core Team, 2020, https://www.R-project.org/).

## Notes

### Competing Interest Statement

The authors have declared no competing interest.

