## Supplementary figures and images for "Protocerebral bridge neurons that regulate sleep in *Drosophila melanogaster*"

### PB-sleep_Fig2_supplement1

#### Figure 2-figure supplement 1

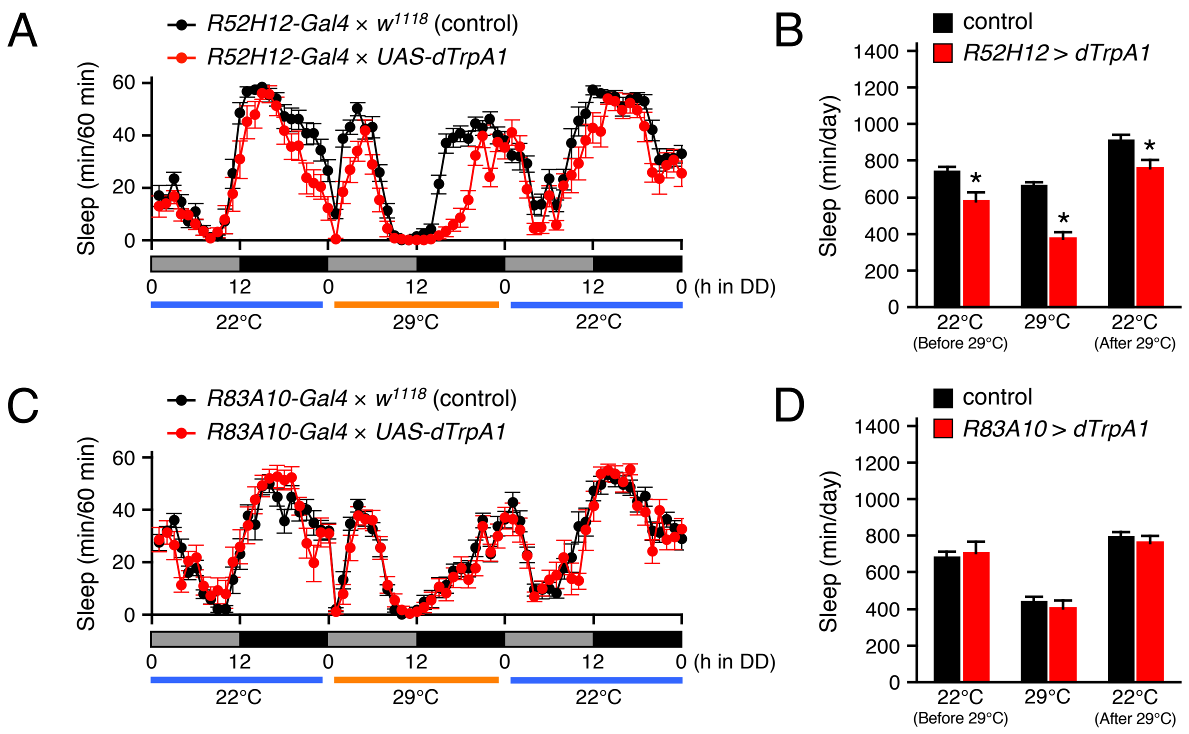

### PB-sleep_Fig2_supplement2

#### Figure 2-figure supplement 2

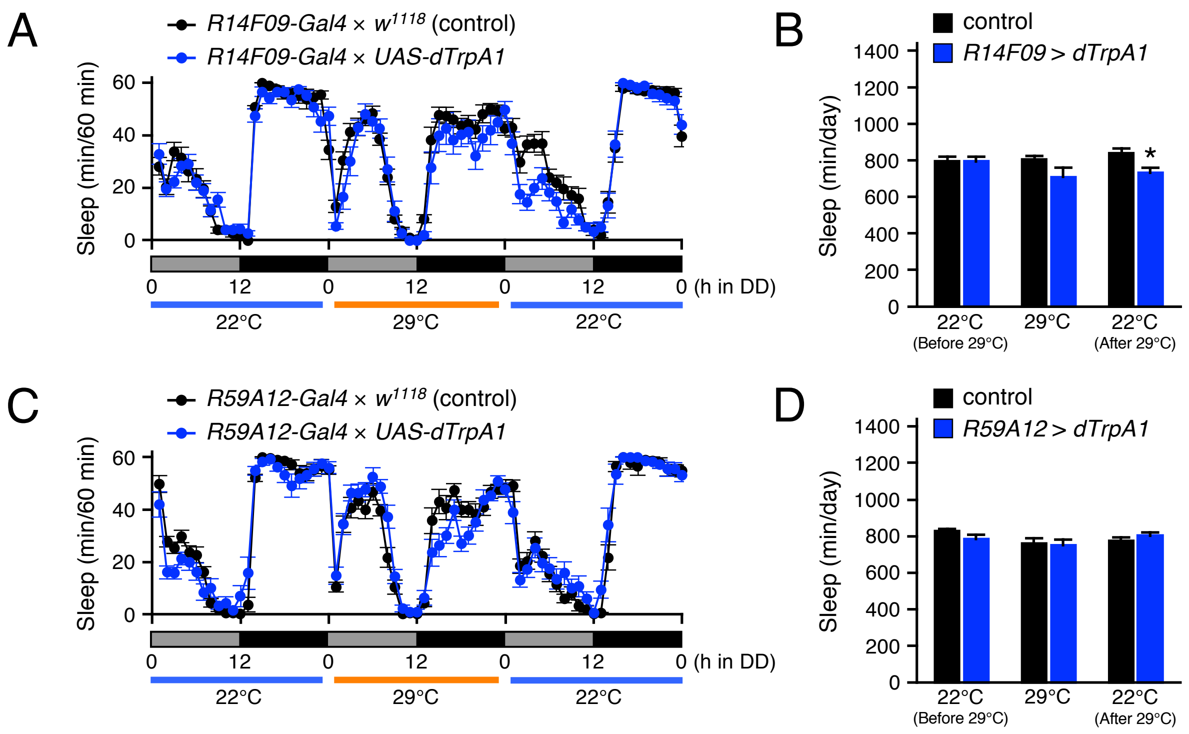

### PB-sleep_Fig7_supplement1

#### Figure 7-figure supplement 1

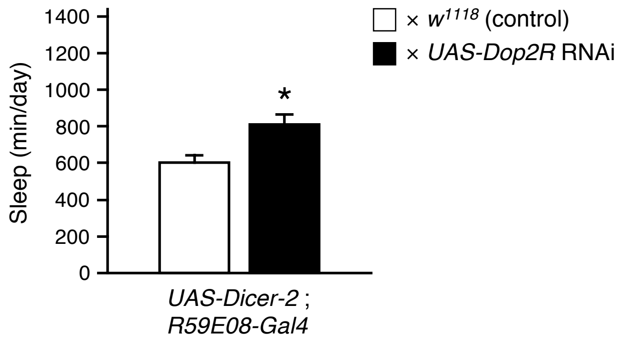

### PB-sleep_Fig7_supplement2

Figure 7-figure supplement 2

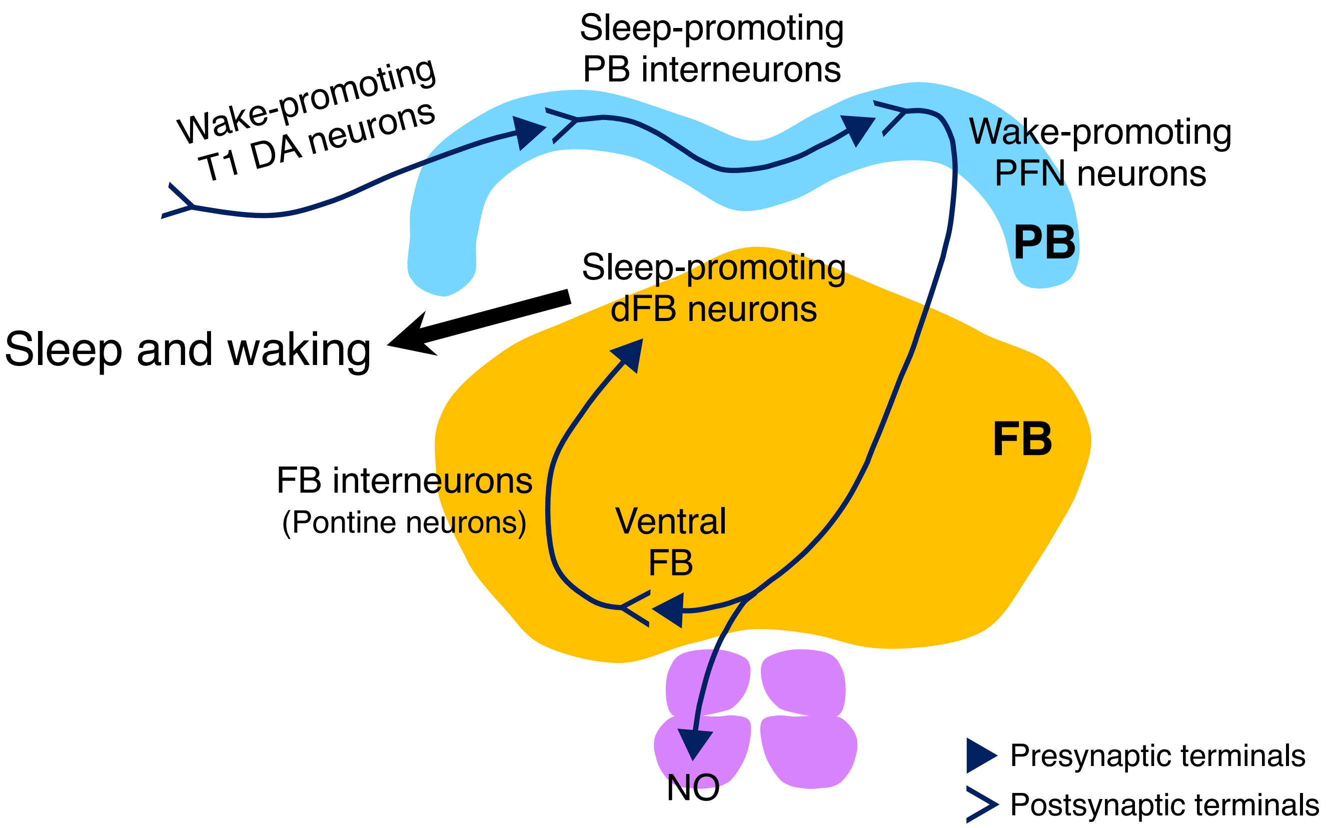
